# Insect immune resolution with EpOME/DiHOME and its dysregulation by their analogs leading to pathogen hypersensitivity

**DOI:** 10.1101/2023.07.07.548078

**Authors:** Md Tafim Hossain Hrithrik, Dong-Hee Lee, Nalin Singh, Anders Vik, Bruce D. Hammock, Yonggyun Kim

## Abstract

Epoxyoctadecamonoenoic acids (EpOMEs) are epoxide derivatives of linoleic acid (9,12-octadecadienoic acid: LA). They are metabolized into dihydroxyoctadecamonoenoic acids (DiHOMEs) in mammals. Unlike in mammals where they act as adipokines or lipokines, EpOMEs act as immunosuppressants in insects. However, the functional link between EpOMEs and pro-immune mediators such as PGE_2_ is not known. In addition, the physiological significance of DiHOMEs is not clear in insects. This study analyzed the physiological role of these C18 oxylipins using a lepidopteran insect pest, *Spodoptera exigua*. Immune challenge of *S. exigua* rapidly upregulated the expression of the phospholipase A_2_ gene to trigger C20 oxylipin biosynthesis, followed by the upregulation of genes encoding EpOME synthase (*SE51385*) and a soluble epoxide hydrolase (*Se-sEH*). The sequential gene expression resulted in the upregulations of the corresponding gene products such as PGE_2_, EpOMEs, and DiHOMEs. Interestingly, only PGE_2_ injection without the immune challenge significantly upregulated the gene expression of *SE51825* and *Se-sEH*. The elevated levels of EpOMEs acted as immunosuppressants by inhibiting cellular and humoral immune responses induced by the bacterial challenge, in which 12,13-EpOME was more potent than 9,10-EpOME. However, DiHOMEs did not inhibit the cellular immune responses but upregulated the expression of antimicrobial peptides selectively suppressed by EpOMEs. The negative regulation of insect immunity by EpOMEs and their inactive DiHOMEs were further validated by synthetic analogs of the linoleate epoxide and corresponding diol. Furthermore, inhibitors specific to Se-sEH used to prevent EpOME degradation significantly suppressed the immune responses. The data suggest a physiological role of C18 oxylipins in resolving insect immune response. Any immune dysregulation induced by EpOME analogs or sEH inhibitors significantly enhanced insect susceptibility to the entomopathogen, *Bacillus thuringiensis*.

**Author summary:** Upon immune challenge, recognition signal triggers insect immunity to remove the pathogens by cellular and humoral responses. Various immune mediators propagate the immune signals to nearby tissues, in which polyunsaturated fatty acid (PUFA) derivatives play crucial roles. However, little was known on how the insects terminate the activated immune responses after pathogen neutralization. Interestingly, C20 PUFA was detected at the early infection stage and later C18 PUFAs were induced in a lepidopteran insect, *Spodoptera exigua*. This study showed the role of epoxyoctadecamonoenoic acids (EpOMEs) in the immune resolution at the late infection stage to quench the excessive and unnecessary immune responses. In contrast, dihydroxy-octadecamonoenoates (DiHOMEs) were the hydrolyzed and inactive forms of EpOMEs. The hydrolysis is catalyzed by soluble epoxide hydrolase (sEH). Inhibitors specific to sEH mimicked the immunosuppression induced by EpOMEs. Furthermore, the inhibitor treatments significantly enhanced the bacterial virulence of *Bacillus thuringiensis* against *S. exigua*. This study proposes a negative control of the immune responses using EpOME/DiHOME in insects.

## 1. Introduction

Epoxyoctadecamonoenoic acids (EpOMEs) including coronaric acid (9,10-EpOME) and vernolic acid (12,13-EpOME) are derived from linoleic acid (LA) by the catalytic activities of specific cytochrome P450 monooxygenases (CYPs) belonging to CYP2C and CYP2J subfamilies in mammalian neutrophils and macrophages [1]. EpOMEs mediate relevant inflammatory response upon pathogen infection [2]. However, excessive EpOMEs are associated with acute respiratory distress syndrome in humans and in rodent models presumably mediated via corresponding diols, 9,10-dihydroxy-octadecamonoenoate (9,10-DiHOME) and 12,13-dihydroxy-octadecamonoenoate (12,13-DiHOME) [3–4]. The DiHOMEs can be detoxified using selective inhibitors of the mammalian soluble epoxide hydrolase (sEH, [5–6]).

In insects, these EpOMEs and DiHOMEs have been speculated to play a crucial role in immune response [7]. Indeed, they were detected in all the developmental stages of the mosquito, *Culex quinquefasciatus* [7]. Feeding sEH inhibitor increased EpOME levels in the mosquito midgut and reduced the total number of bacteria in the lumen, suggesting that EpOMEs were associated with insect immunity [8]. The role of EpOMEs in insect immune response was further analyzed in the immune-challenged larvae of a lepidopteran insect, *Spodoptera exigua*, which exhibited relatively high levels of 941.8 pg/g of 9,10-EpOME and 2,198.3 pg/g of 12,13-EpOME [9]. In the insect species, a specific cytochrome P450 enzyme (CYP, *SE51385*) and a soluble epoxide hydrolase (*Se-sEH*) mostly catalyzed the production and hydrolysis of EpOMEs, respectively. These two regioisomers of EpOMEs suppress the cellular and humoral immune responses. Their immunosuppressive effects were further supported by treatment with urea-based sEH inhibitors, which inhibited the cellular and humoral immune responses of *S. exigua* probably by increasing the levels of EpOMEs in response to immune challenge. Thus, EpOMEs facilitate anti-inflammatory response in the insect. However, little is known regarding the role of EpOMEs and their association with pro-inflammatory agents.

DiHOMEs are also produced in neutrophils in humans following bacterial challenge [10–11]. Although DiHOMEs are the metabolites of EpOMEs, they act as mediators of neutrophil chemotaxis [12]. High concentrations of DiHOMEs under physiological stress such as following severe skin burn or COVID infection can lead to toxicity by acting as isoleukotoxins, resulting in severe dysfunction of the innate immune responses [6]. DiHOMEs also suppress excessive production of reactive oxygen species in neutrophils [4]. However, studies investigating the physiological role of DiHOMEs in insects have yet to be reported.

This study investigated the physiological role of C18 oxylipins in attenuating the immune response in insects by analyzing the functional relationship between EpOMEs and pro-immune mediators as well as EpOME metabolism. The physiological roles of the resulting DiHOMEs in insect immunity were also investigated. The physiological role of C18 oxylipins in immune resolution was determined by analyzing the effects of DiHOME analogs or sEH inhibitors on dysregulation of the immune responses and susceptibility to an insect pathogen.

## 2. Results

### 2.1 Sequential synthesis of PGE_2_, EpOMEs, and DiHOMEs in immune-challenged larvae of *S. exigua*

Bacterial challenge induced the expressions of *PLA_2_*, EpOME synthase (*SE51385*), and *Se-sEH* genes in *S. exigua* larvae (Fig 1A). However, their expression profiles differed (*F* = 3.96; df = 2, 9; *P* = 0.0009), and the maximum levels were detected during peak times. The maximal expression peak of *PLA_2_* was observed at 1 h after the bacterial challenge while the peaks of *SE51385* and *Se-sEH* were detected 4 h post-challenge. In addition, *SE51385* was expressed earlier (*F* = 5.70; df = 1, 9; *P* < 0.0001) than *Se-sEH*. Interestingly, only PGE_2_ injection induced the gene expression of *SE51385* and *Se-sEH* (Fig 1B).

**Fig 1.**
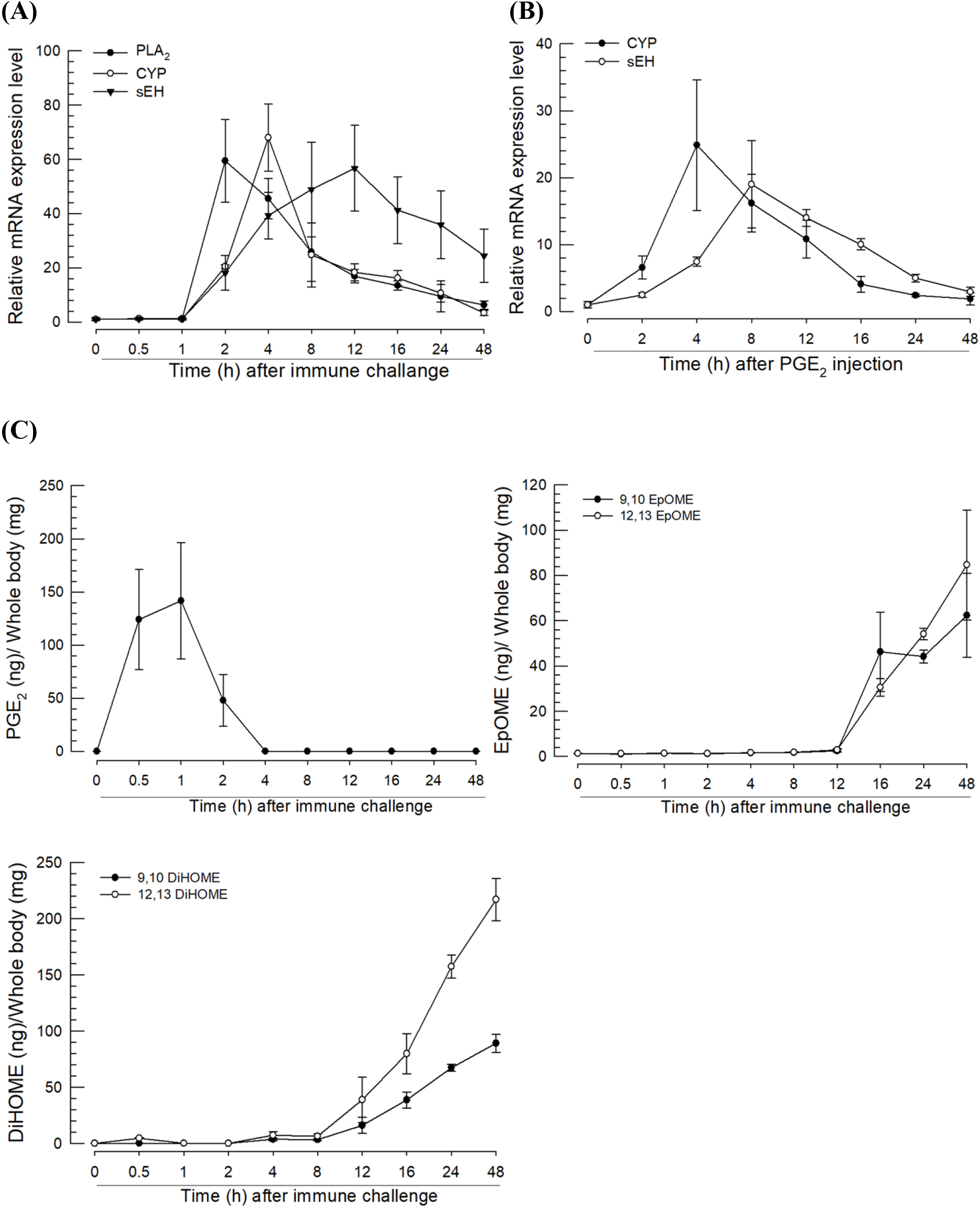
Expression profiles of phospholipase A_2_ (*PLA_2_*, GenBank accession number: MH061374.1), EpOME synthase (*CYP*, MT375603.1), and soluble epoxide hydrolase (*sEH*, MT375604.1). (**A**) Inducible expressions of three genes in response to bacterial challenge. *E. coli* was injected into the larval proleg (1.73 × 10^3^ cells/larva). After 8 h incubation, the total extracted RNA from the fat body was used to determine the mRNA expression level. Different levels of expression in RT-qPCR samples were normalized using the expression levels of a ribosomal protein, *RL32*. (**B**) Inducible expressions of *CYP* and *sEH* after PGE_2_ injection. PGE_2_ was injected into each larva at a dose of 1 µg. (**C**) Quantification of PGE_2_ and EpOMEs/DiHOMEs in the whole body without the gut in *S. exigua* larvae. The samples were collected from the challenged larvae, in which L4 larvae were challenged with *E. coli* (1.73 × 10^3^ cells/larva). After injection, the total body except the gut was collected at different time points and used to measure PGE_2_, EpOME, and DiHOME levels using LC–MS/MS. Each assessment was replicated with three independent samples.

LC-MS/MS was used to analyze the concentrations of PGE_2_, EpOMEs, and DiHOMEs in immune-challenged larvae at different time points (Fig 1C). To avoid exogenous dietary contamination, the entire gut was removed from the whole body preparation before extracting the oxylipins. The immune challenge significantly increased the levels of PGE_2_ and EpOMEs/DiHOMEs. The induced levels of PGE_2_ were detected earlier (0.5 ∼ 2 h) than those (> 12 h) of EpOMEs/DiHOMEs. The induced levels of the two EpOMEs did not differ significantly (*F* = 0.26; df = 1, 9; *P* = 0.6138). However, the concentrations of the two DiHOMEs showed statistically significant differences (*F* = 17.46; df = 1, 9; *P* < 0.0001) following induction, and 12,13-DiHOME was detected at higher levels than 9,10-DiHOME.

### 2.2 EpOMEs, but not DiHOMEs, suppress immune response

To investigate the role of EpOMEs/DiHOMEs in insect immunity, hemocyte-spreading behavior was assessed by adding them to the hemocyte suspension (left panel in Fig 2A). Control hemocytes spread via cytoplasmic extensions along with increased levels of F-actin. However, the addition of 12,13-EpOME inhibited hemocyte spread in a dose-dependent manner (Fig 2B). Treatment with LA, its biosynthetic precursor, also slightly suppressed the hemocyte spread. However, 12,13-DiHOME did not suppress even at the highest test concentration. Compared with 9,10-EpOME, 12,13-EpOME exhibited higher suppression of hemocyte-spreading behavior. However, none of the regioisomeric DiHOMEs inhibited (*F* = 2.92; df = 1, 4; *P* = 0.1625) the spread of hemocytes.

**Fig 2.**
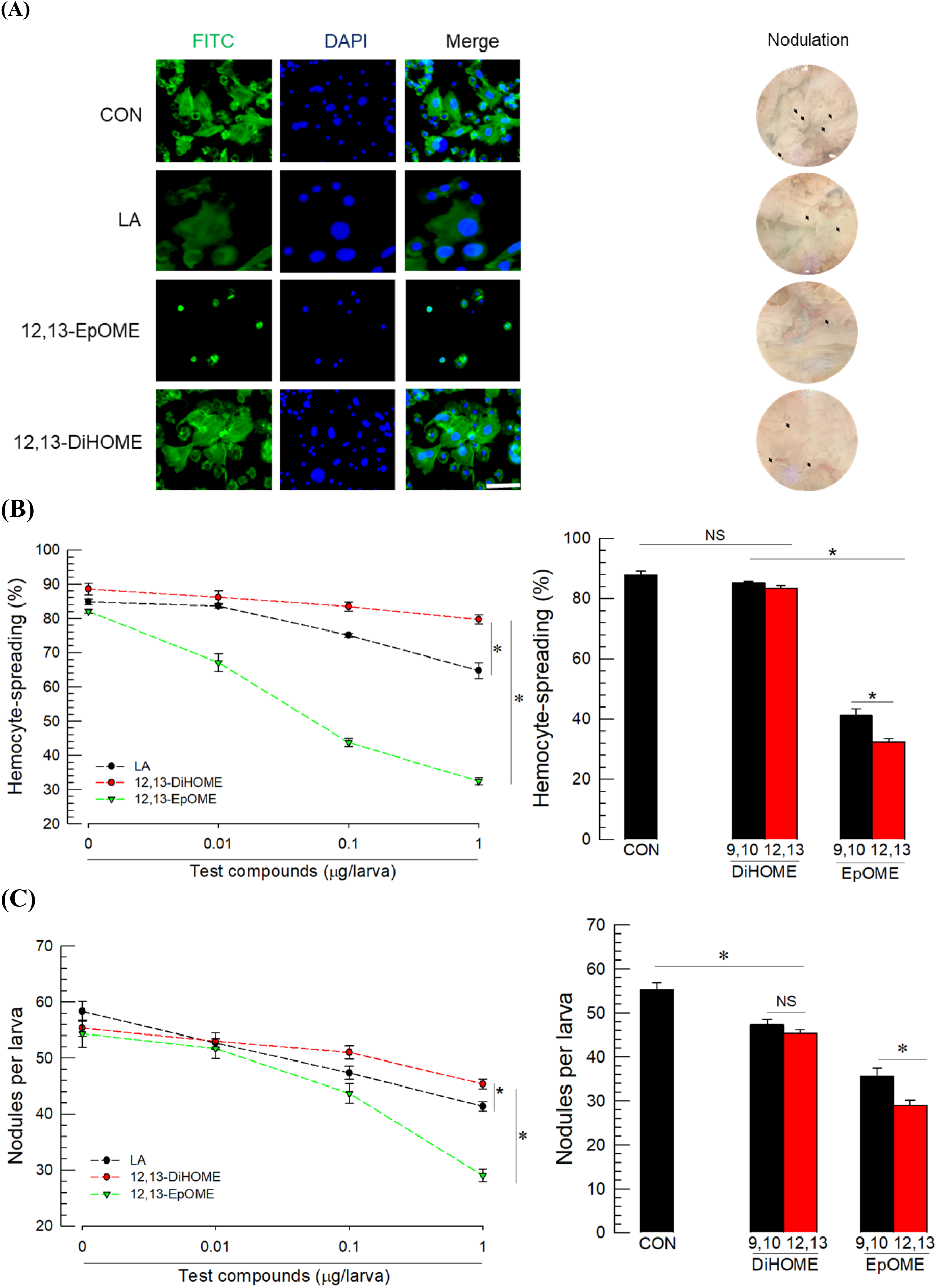
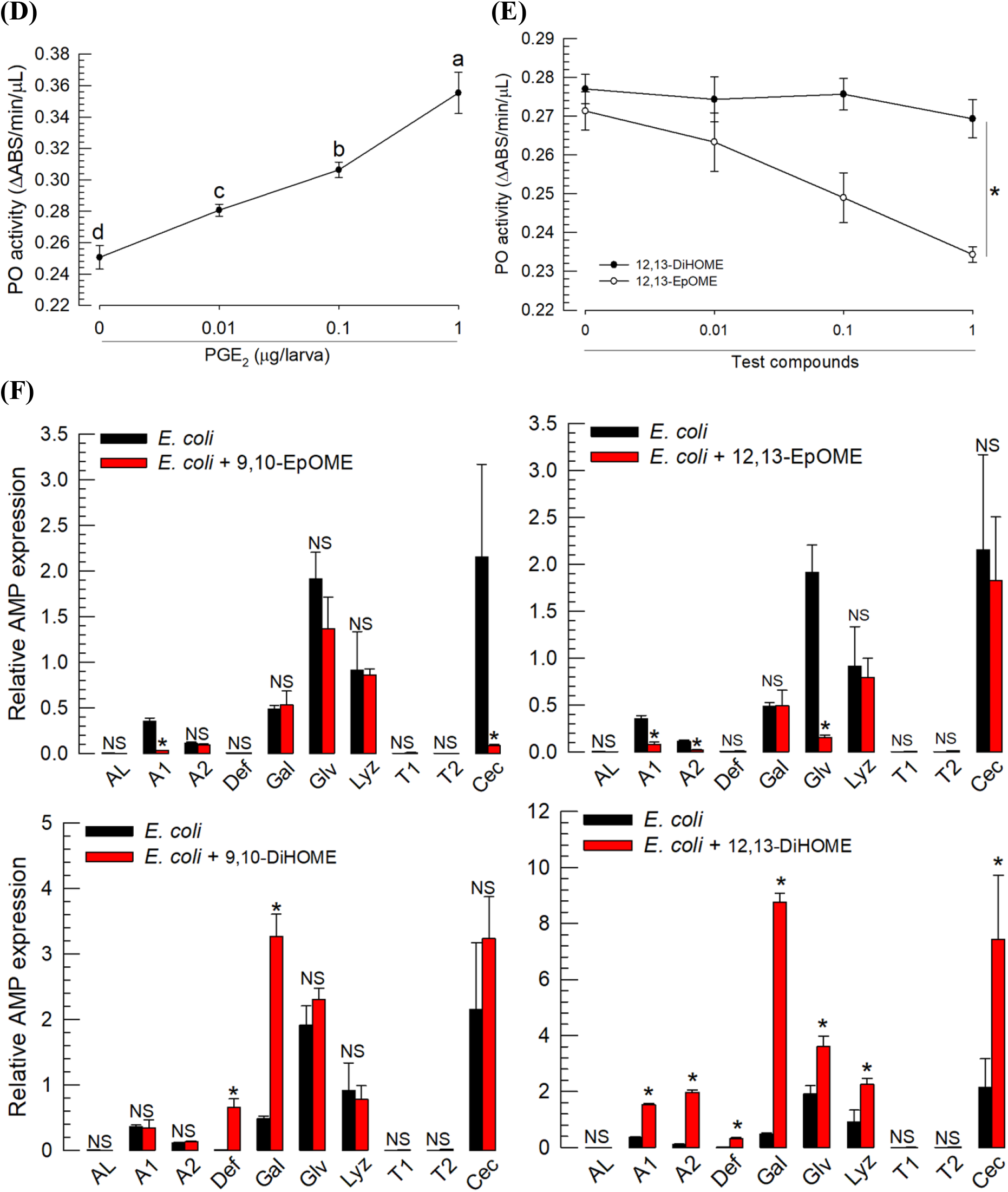
Contrasting effects of EpOMEs and DiHOMEs on immune responses of *S. exigua*. (**A**) Hemocyte-spreading behavior (left) and nodule formation (right) after exposure to test compounds including linoleic acid (LA). Hemocytes were stained with FITC against F-actin and DAPI against nucleus. Nodules are indicated by arrows. The scale bar represents 10 µm. (**B**) Effect of different doses of test compounds on hemocyte-spreading behavior. The cellular spread was scored by counting the number of cells exhibiting F-actin growth out of cell boundary among 100 randomly selected cells. The right panel compares the effects of a single dose (1 µg) of DiHOME and EpOME in each regioisomer on hemocyte spread. (**C**) Effect of different doses of test compounds on nodulation. Nodule formation was determined by dissection at 8 h after injection with *E. coli* (1.73×10^3^ cells/larva) and/or test compounds. For control (‘CON’) treatment, only *E. coli* was injected. Right panel shows a comparison of nodule formation after exposure to a single dose as described above. (**D**) Upregulation of phenol oxidase (PO) activity at 8 h after PGE_2_ (1 µg per larva) treatment. (**E**) Downregulation of PO activity with test compounds (µg) injected along with the bacteria. (**F**) Effects of EpOMEs and DiHOMEs on the expression of 10 antimicrobial peptide (AMP) genes: apolipophorin III (*AL*), defensin (*Def*), gallerimycin (*Gal*), gloverin (*Glv*), lysozyme (*Lyz*), transferrin I (*T1*), transferrin II (*T2*), attacin 1 (*A1*), attacin 2 (*A2*), and cecropin (*Cec*). AMP gene expression was determined via RT-qPCR at 8 h after injection with *E. coli* (1.73×10^3^ cells/larva) and/or test compounds. Each treatment was replicated three times with individual tissue preparations. ‘NS’ stands for no significant difference between two treatments. Asterisks or different letters above standard error bars indicate significant differences among means with a type I error of 0.05 (LSD test).

Nodule formation (right panel in Fig 2A) was assessed to test the effect of oxylipin on the cellular immune response of *S. exigua*. Bacterial challenge resulted in the formation of about 55 nodules per larva in the hemocoel (Fig 2C). Both regioisomeric EpOMEs significantly suppressed the nodule formation, although 12,13-EpOME was more potent than 9,10-EpOME and exhibited dose-dependent inhibition. LA also suppressed the nodule formation but was less potent. However, none of the DiHOMEs exhibited any inhibitory activity (*F* = 1.80; df = 1, 4; *P* = 0.2508) against the cellular immune response.

Melanization occurs in cellular immune responses via upregulation of PO activity in *S. exigua* [13]. PGE_2_ treatment induced the upregulation of PO activity in a dose-dependent manner (Fig 2D). However, 12,13-EpOME significantly suppressed the PO activity, whereas 12,13-DiHOME did not alter PO activity (Fig 2E)

To test the influence of DiHOMEs on the humoral immune responses, 10 AMP genes induced by immune challenge were assessed (Fig 2F). Both EpOME regioisomers suppressed the expression of a few AMPs following immune challenge. The regioisomer 9,10-EpOME suppressed *attacin 1* and *cecropin* genes while 12,13-EpOME suppressed *attacin 1*, *attacin 2*, and *gloverin*. Interestingly, both DiHOMEs upregulated the expression of the four AMPs suppressed by the EpOMEs. In addition, 9,10-DiHOMEs upregulated the expression of *defensin* and *gallerimycin* genes while 12,13-DiHOME upregulated *gloverin* and *lysozyme* genes.

### 2.3 EpOMEs exhibit hemolytic activity at low concentrations while DiHOMEs do not

The analysis of hemocyte behavior in response to 12,13-EpOME treatment revealed cellular blebbing. (Fig 3A). However, 12,13-DiHOME treatment had limited effect on cell blebbing in hemocytes. The cell blebbing was further analyzed under different concentrations of EpOMEs/DiHOMEs (Fig 3B). Both EpOMEs significantly induced cell blebbing: 12,13-EpOME was more potent than 9,10-EpOME and exhibited hemolytic activity in a dose-dependent manner. LA also induced cell blebbing but its activity was substantially lower than that of EpOMEs. However, 9,10-DiHOME did not exhibit hemolytic activity. In contrast, 12,13-DiHOME was slightly hemolytic but significantly weaker than LA.

**Fig 3.**
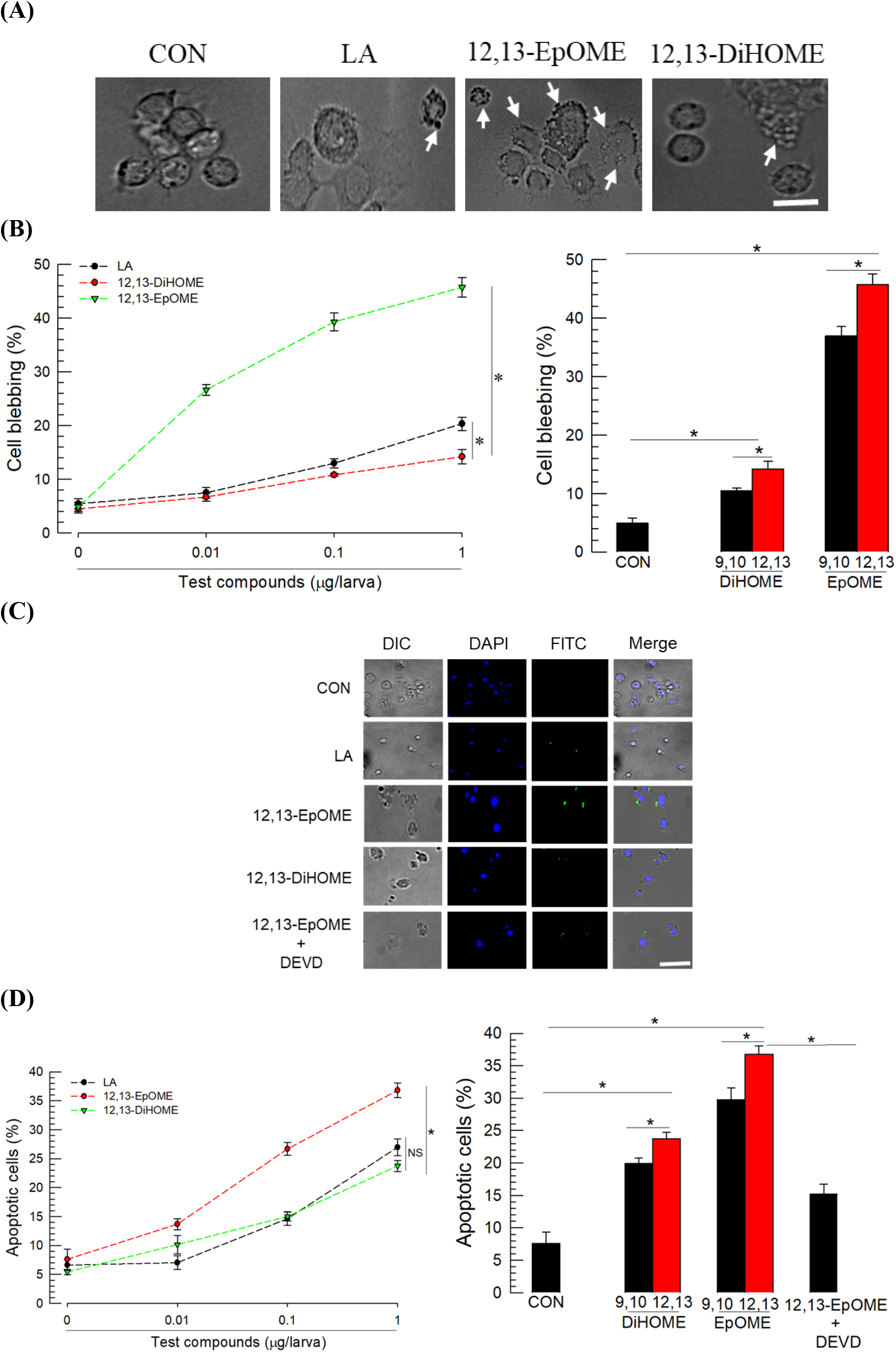
Contrasting cytotoxic effects of EpOMEs and DiHOMEs against *S. exigua* hemocytes. (**A**) Cytotoxicity manifested by hemocyte membrane blebbing (arrows) after exposure to test compounds including linoleic acid (LA). The scale bar represents 10 µm. (**B**) Effect of different doses of test compounds on cellular blebbing. The right panel compares the cytotoxic effects of a single dose (1 µg) of DiHOME and EpOME in each of regioisomers. (**C**) Apoptotic effects characterized by DNA fragmentation (stained by FITC against BrdU) of hemocytes after exposure to test compounds including DEVD (a caspase inhibitor) using TUNEL assay. Nucleus was stained with DAPI. Cells were visualized under differential interference contrast (DIC). (**D**) Apoptotic effects of different doses of test compounds. The right panel compares the apoptotic effects of a single dose (1 µg) of DiHOME and EpOME in each regioisomer. Apoptosis was scored by counting the number of FITC-stained cells among 100 randomly selected cells. Each treatment was replicated three times with individual tissue preparations. ‘NS’ stands for no significant difference between two treatments. Asterisks or different letters above standard error bars indicate significant differences among means with a type I error of 0.05 (LSD test).

The cytotoxicity manifested by cellular blebbing was further analyzed by TUNEL assays to determine whether the hemolytic activity was caused by apoptosis (Fig 3C). Incubation of hemocytes with 12,13-EpOME led to DNA fragmentation, which was observed by DNA end labeling with BrdU. The DNA fragmentation increased in a dose-dependent manner following incubation with 12,13-EpOME. Treatment with 9,10-EpOME also induced DNA fragmentation but less than in 12,13-EpOME treatment (Fig 3D). Treatment with Ac-DEVD-AMC and Caspase-3 inhibitor (Ac-DEVD-CHO) significantly rescued the DNA fragmentation induced by 12,13-EpOME. LA also induced the DNA fragmentation but its activity was substantially lower than that of EpOMEs. Both DiHOMEs also exhibited apoptotic activity at high concentrations but their activities were significantly lower than those of EpOMEs.

### 2.4 Effect of other LA metabolites on immune response

To analyze the physiological roles of EpOMEs and their hydrolytic products (DiHOMEs) in insects, different LA metabolites were tested by preparing two epoxide-containing diEpOMEs and two different tetrahydroxy metabolites (THOME and THF-diols) (S2A Fig). As expected, diEpOMEs similar to EpOMEs strongly suppressed cellular immune response such as hemocyte-spreading behavior (Fig 4A) and nodule formation (Fig 4B). They also inhibited PO activation (Fig 4C). The diEpOME significantly suppressed the induction of some AMP genes (Fig 4D). In addition, diEpOME showed high levels of cytotoxicity against hemocytes by inducing apoptosis (Fig. 4E and 4F). In contrast, THOME, similar to DiHOMEs, did not inhibit the cellular immune response upon immune challenge (Fig 4A and 4D). Similar to DiHOMEs, it induced humoral immune response by further upregulating the gene expression of some AMPs induced by the bacterial infection (Fig 4D). Interestingly, THF-diols slightly retained the inhibitory activities against cellular and humoral immune responses but were substantially weaker than those of the epoxide metabolites. Both THOME and the THF-diol were less cytotoxic to hemocytes compared with diEpOME (Fig 4E and 4F).

**Fig 4.**
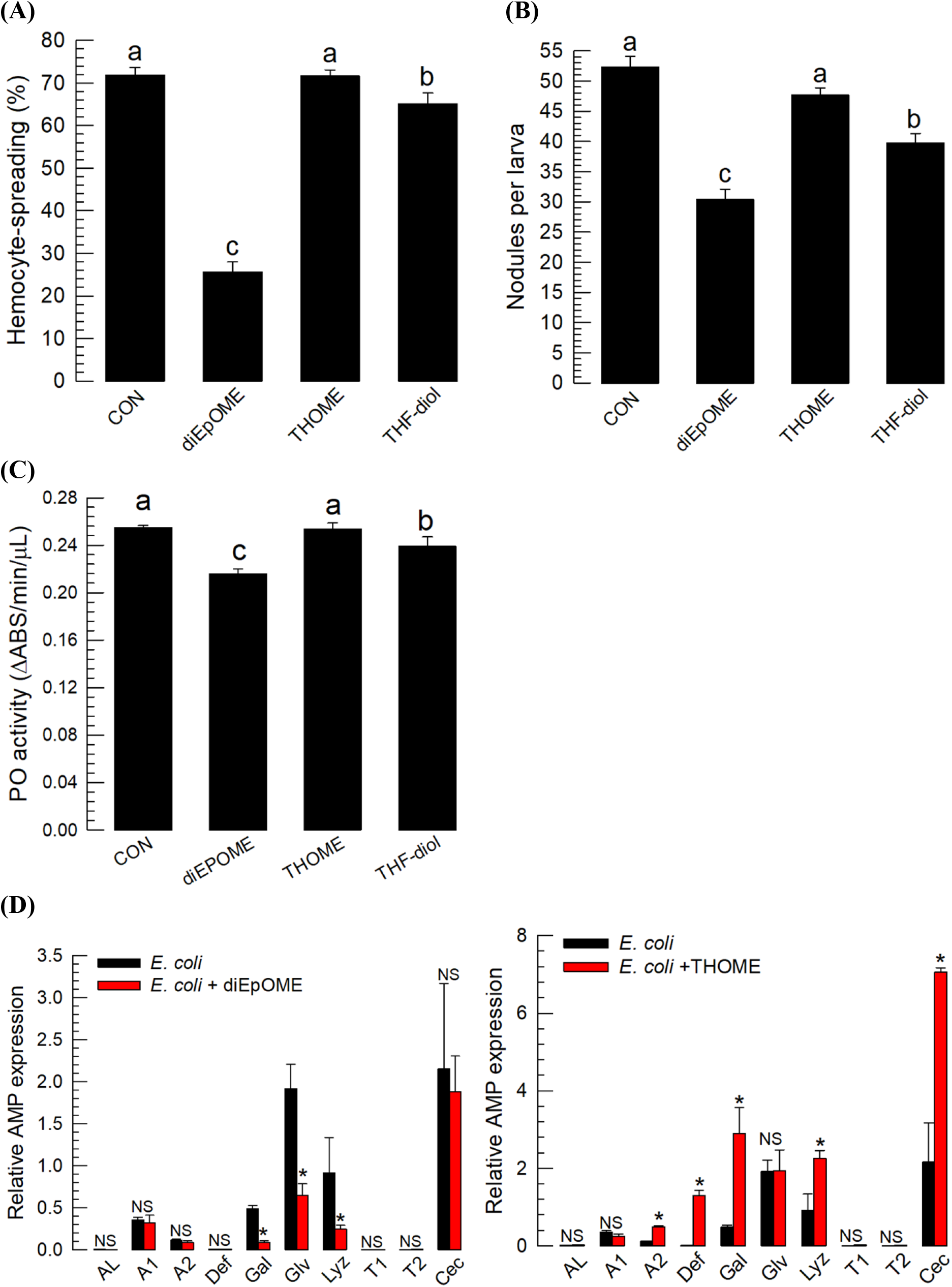
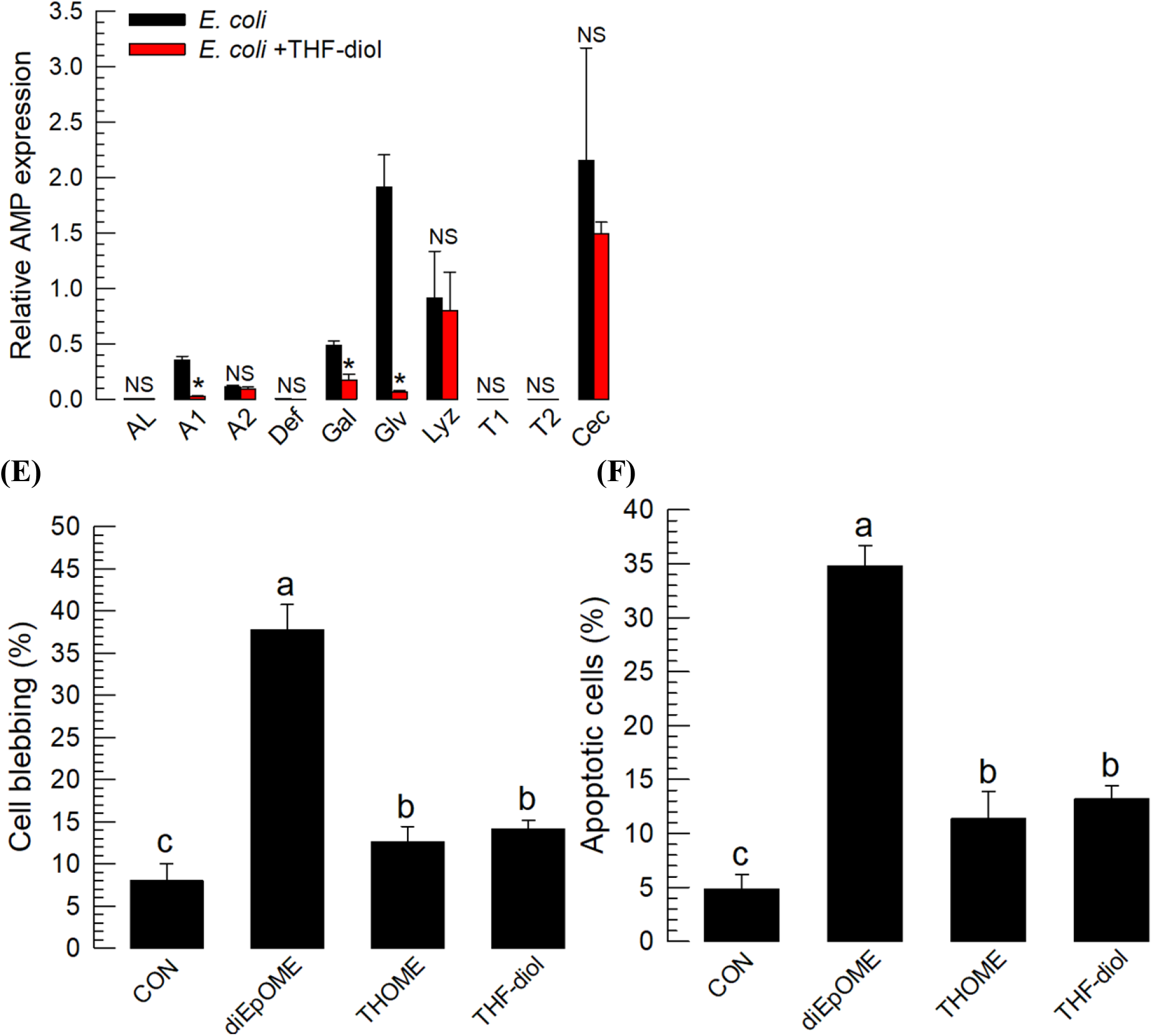
Effects of three different linoleic acid (LA) metabolites (diEpOME, THOME, and THF-diol, see Fig. S2A) on immune responses of *S. exigua*. (**A**) Effects of three LA metabolites (1 µg/larva) on hemocyte-spreading behavior. The cellular spread was determined by counting the cells exhibiting F-actin growth (stained by FITC) out of cell boundary among 100 randomly selected cells. (**B**) The effect (1 µg/larva) on nodule formation at 8 h after bacterial infection (1.73 × 10^3^ cells/larva) is shown. (**C**) The effect (1 µg/larva) on phenol oxidase (PO) activity at 8 h after bacterial infection (1.73×10^3^ cells/larva) is shown. (**D**) Effect (1 µg/larva) on the expression of 10 antimicrobial peptide (AMP) genes: apolipophorin III (*AL*), defensin (*Def*), gallerimycin (*Gal*), gloverin (*Glv*), lysozyme (*Lyz*), transferrin I (*T1*), transferrin II (*T2*), attacin 1 (*A1*), attacin 2 (*A2*), and cecropin (*Cec*). Expression of AMP genes via RT-qPCR at 8 h after injection with *E. coli* (1.73×10^3^ cells/larva) and/or test compounds.. Each treatment was replicated three times with individual tissue preparations. The cytotoxic effects of each treatment (1 µg/larva) on hemocytes was measured based on cellular blebbing (**E**) and apoptosis using TUNEL assay (**F**). Each cytotoxicity test was performed by counting the number of cell blebs or FITC-stained cells for apoptosis among 100 randomly chosen cells. Each treatment was replicated three times. ‘NS’ stands for no significant difference between two treatments. Asterisks or different letters above standard error bars indicate significant differences among means with a type I error of 0.05 (LSD test).

### 2.5 Effect of EpOME alkoxides on immune response

The immunosuppressive activities of EpOMEs were further analyzed using stable alkoxide derivatives (S2B Fig). Methoxy (A839, A843), ethoxy (A840, A844), propoxy (A841, A845), and *iso*-propoxy (A842, A846) ethers were used as bioisosteres for the epoxide in the EpOMEs. Each alkoxide derivative was prepared and tested as a regioisomeric mixture of 9-, 10-, 12- and 13-alkoxyoctadecenoic acids or methyl esters. Among the eight alkoxide derivatives, the methyl ester A841 was the most potent suppressor of cellular immune response in a dose-dependent manner (Fig 5A-5C). The analysis of AMP expression revealed the relative potency of A841 in suppressing the induction of AMP genes (Fig 5D) compared with those of both EpOMEs (Fig 2F). A841, similar to 12,13-EpOME, was also cytotoxic to hemocytes and induced cellular blebbing (Fig 5E) and apoptosis (Fig 5F).

**Fig 5.**
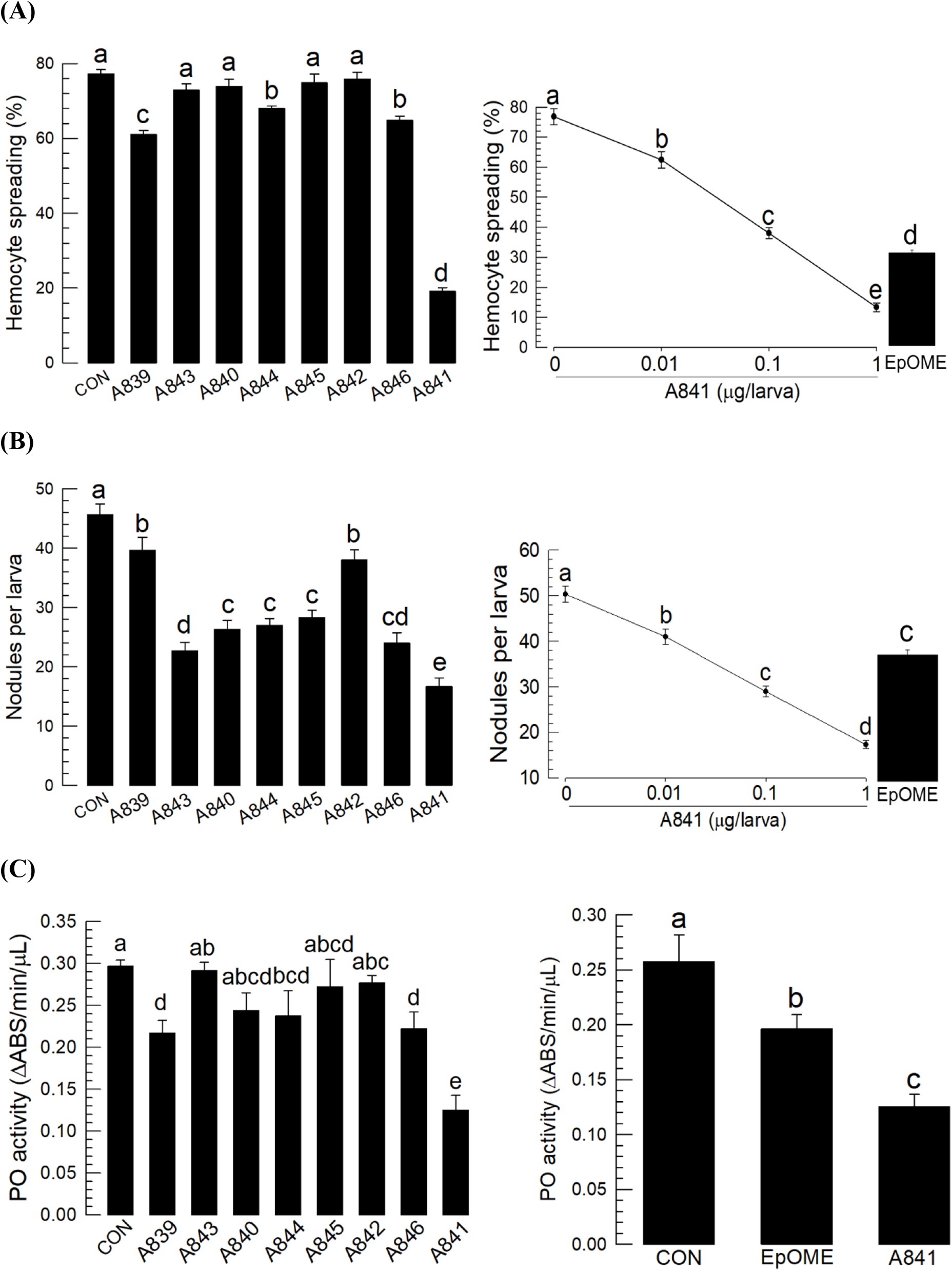
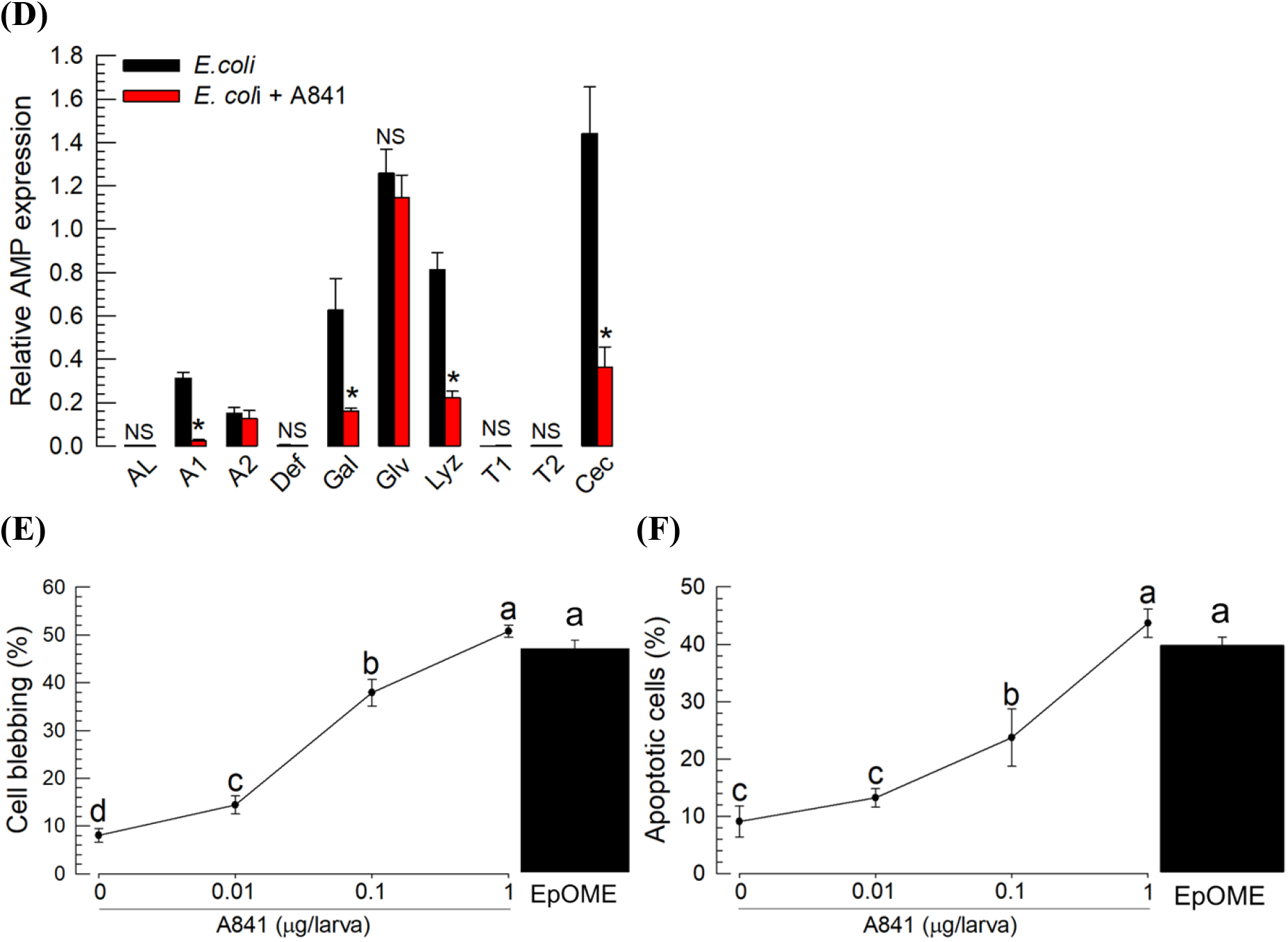
Effects of eight different EpOME alkoxides (see S2B Fig) on immune responses of *S. exigua*. (**A**) Effects of EpOME alkoxides (1 µg/larva) on hemocyte-spreading behavior. The cellular spread was determined by counting the cells exhibiting F-actin growth (stained by FITC) out of cell boundary among 100 randomly selected cells. Right panel shows the inhibitory activities of different doses of A841 on hemocyte-spreading behavior compared with 12,13-EpOME treatment (1 µg/larva) (**B**) The effect of treatment (1 µg/larva) on nodule formation at 8 h after bacterial infection (1.73×10^3^ cells/larva). Right panel shows the inhibitory activities of different doses of A841 on nodulation compared with the effects of 12,13-EpOME (1 µg/larva) (**C**) The effect of treatment (1 µg/larva) on phenol oxidase (PO) activity at 8 h after bacterial infection (1.73×10^3^ cells/larva). Right panel shows the inhibitory activities of different doses of A841 on PO activity compared with the effects of 12,13-EpOME (1 µg/larva). (**D**) Effect of A841 (1 µg/larva) on the expression of 10 antimicrobial peptide (AMP) genes: apolipophorin III (*AL*), defensin (*Def*), gallerimycin (*Gal*), gloverin (*Glv*), lysozyme (*Lyz*), transferrin I (*T1*), transferrin II (*T2*), attacin 1 (*A1*), attacin 2 (*A2*), and cecropin (*Cec*). The expression of AMP genes via RT-qPCR was evaluated at 8 h after injection with *E. coli* (1.73 × 10^3^ cells/larva) and/or test compounds. Each treatment was replicated three times with individual tissue preparations. Cytotoxic effects of A841 (1 µg/larva) on hemocytes were measured in terms of cellular blebbing (**E**) and apoptosis using TUNEL assay (**F**). Each cytotoxicity test was performed by counting the number of cell blebs or FITC-stained cells (for apoptosis) among 100 randomly chosen cells. Each treatment was replicated three times. ‘NS’ stands for no significant difference between two treatments. Asterisks or different letters above standard error bars indicate significant difference among means with a type I error of 0.05 (LSD test).

### 2.6 Effect of sEH inhibitors on immune responses

The immunosuppressive activity of EpOMEs suggested that sEH inhibitors suppress the immune responses of *S. exigua* by elevating the endogenous EpOME levels following the immune challenge. Six urea-based sEH inhibitors were assessed to test the hypothesis. All sEH inhibitors significantly inhibited the cellular (Fig 6A-6C) or humoral responses (Fig 6D). The sEH inhibitors AUDA and CUDA were most potent inhibitors. These two sEH inhibitors were also highly cytotoxic to hemocytes and induced cellular blebbing (Fig 6E) and apoptosis (Fig 6F).

**Fig 6.**
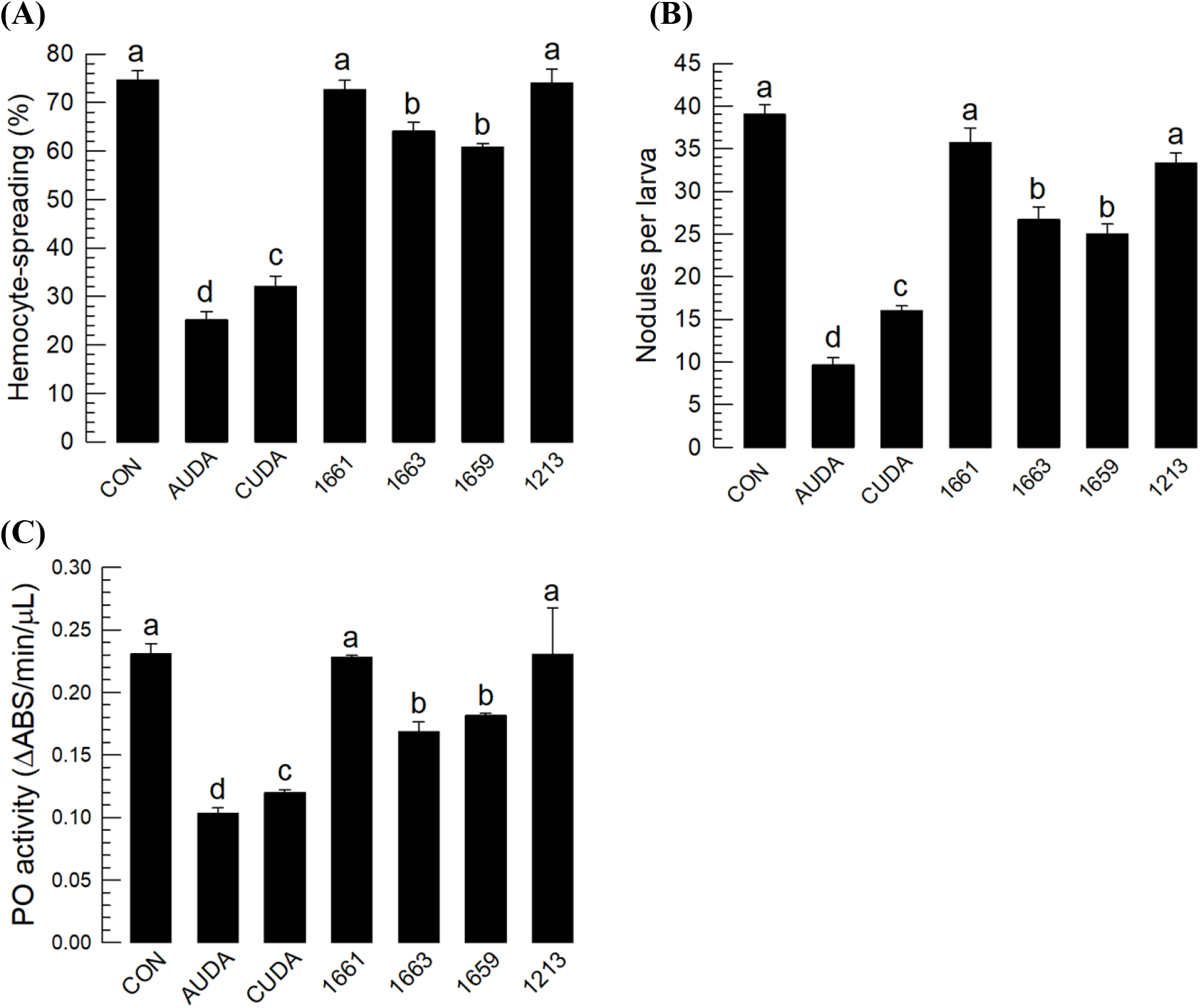
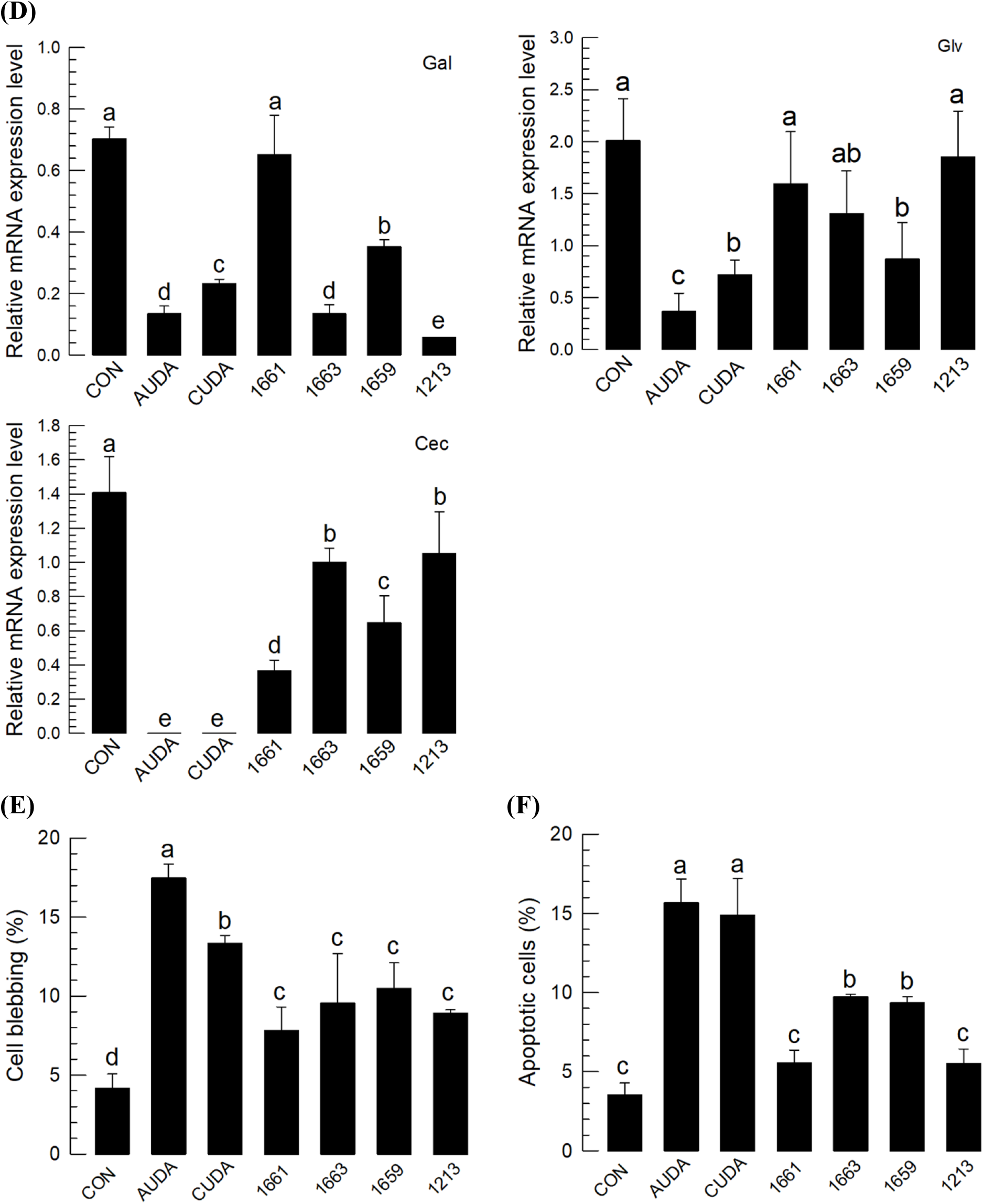
Effects of six different sEH inhibitors (see S2C Fig) on immune responses of *S. exigua*. (**A**) Effects of the inhibitors (1 µg/larva) on hemocyte-spreading behavior. The cellular spread was determined by counting the number of cells exhibiting F-actin growth (stained by FITC) out of cell boundary among 100 randomly selected cells. (**B**) Their effect of inhibitors (1 µg/larva) on nodule formation at 8 h after bacterial infection (1.73 × 10^3^ cells/larva). (**C**) The effect of inhibitors (1 µg/larva) on phenol oxidase (PO) activity at 8 h after bacterial infection (1.73 × 10^3^ cells/larva). (**D**) The effect of treatment (1 µg/larva) on the expression of three antimicrobial peptide (AMP) genes: gallerimycin (*Gal*), gloverin (*Glv*), and cecropin (*Cec*). AMP genes were analyzed by RT-qPCR at 8 h after injection with *E. coli* (1.73 × 10^3^ cells/larva) and/or test compounds. Each treatment was replicated three times with individual tissue preparations. Cytotoxic effects of the inhibitors (1 µg/larva) on hemocytes were measured by counting the number of cellular blebs (**E**) and determining the degree of apoptosis using TUNEL assay (**F**). Each cytotoxicity test was performed by counting the number of cell blebs or FITC-stained cells (for apoptosis) among 100 randomly selected cells. Each treatment was replicated three times. ‘NS’ indicates no significant difference between two treatments. Asterisks or different letters above standard error bars indicate significant differences among means with a type I error of 0.05 (LSD test).

### 2.7 Enhanced Bt virulence following treatment of *S. exigua* with diEpOME or sEH inhibitors

The immunosuppressive and hemolytic activities of EpOME derivatives (Fig 7A) and sEH inhibitors suggested their potential role in enhancing the virulence of entomopathogenic microbes. Each of these compounds showed significant insecticidal activities against *S. exigua* larvae, and diEpOME was more potent than the sEH inhibitor (Fig 7B). To test the enhanced virulence hypothesis, *B. thuringiensis* (= an entomopathogenic bacterium) was treated with each of the potent immunosuppressants (Fig 7C). All three treatments significantly enhanced the bacterial virulence, and AUDA and the EpOME-mimic A841 were the most potent compounds.

**Fig 7.**
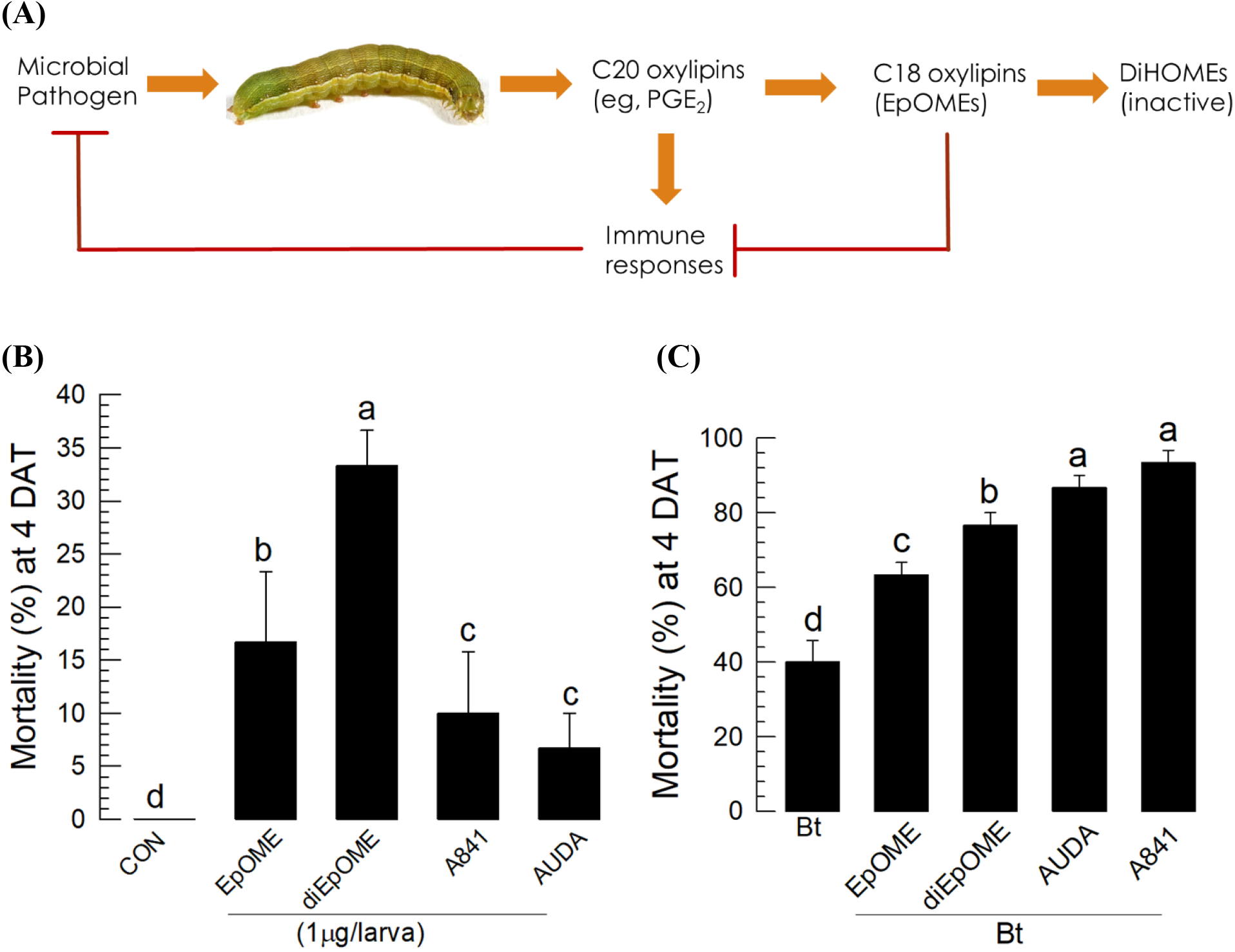
Immunosuppressive activity of EpOMEs and their role in enhancing entomopathogenic virulence against *S. exigua*. (**A**) Diagram illustrating the physiological roles of C18 and C20 oxylipins in modulating insect immunity. (**B**) Insecticidal activities of the four selected compounds (EpOME, diEpOME, A841, and AUDA) against *S. exigua* following injection of hemocoel at a dose of 1 µg/larva to L4 larvae. Controls were injected with DMSO. (**C**) Enhanced virulence of *B. thuringiensis aizawai* (Bt) against *S. exigua*. The larvae injected with the test compounds (1 µg/larva) were fed with cabbage soaked in Bt suspension using a leaf-dipping method. Mortality was assessed at 4 days after treatment (‘DAT’). Ten larvae were used in each treatment and the treatment was performed in triplicate. Different letters above standard error bars indicate significant differences among means with a type I error of 0.05 (LSD test).

## 3. Discussion

Polyunsaturated fatty acids (PUFAs) are fatty acids that contain two or more double bonds in their carbon backbone. The importance of PUFAs as essential fatty acids has been appreciated since the discovery of the health benefits of animal and human diets containing omega-3 and omega-6 fatty acids [1]. In particular, long-chain PUFAs (LC-PUFAs: 20 or more carbon atoms) function as signaling molecules contributing to immunity, development, and reproduction. LC-PUFAs are also rich (7–25%) in aquatic insects, but are rare in terrestrial insects due to evolutionary adaptation to prevent oxidative stress [14]. However, terrestrial insects are rich in LA (18-carbon fatty acid) present in phospholipids (PLs). For example, phospholipids extracted from the fat body of *S. exigua* larvae contained 34.4% LA of the total fatty acids [15]. LA metabolites such as EpOMEs and DiHOMEs are well known for their physiological roles in mammals [1]. However, their physiological role in insects have not received substantial attention. This study assessed the physiological role of LA metabolites in a lepidopteran insect, especially their crucial role in mediating immune responses.

Upon immune challenge, increasing levels of EpOME and DiHOME regioisomers were produced along with PGE_2_ production in *S. exigua*. The immune challenge also sequentially induced the gene expression of PLA_2_, EpOME synthase, and sEH. More interestingly, the injection of PGE_2_ alone induced the gene expression of EpOME synthase and sEH. This suggests that the immune challenge induces PLA_2_ gene expression or directly activates its catalytic activity to synthesize C20 oxylipins including PGE_2_, which activates cellular or humoral immune responses of *S. exigua* [14]. Arachidonic acid and LA in the phospholipids of *S. exigua* facilitate the catalytic activity of PLA_2_ to release the fatty acids, especially LA. LA subsequently undergoes chain elongation and desaturation to produce arachidonic acid, which is used to synthesize various eicosanoids [16]. Subsequent induction of EpOME synthase and sEH enzymes utilizes the free LA to generate EpOME and DiHOME isomers presumably to shut down the induced immune response. The relative fatty acid abundance triggers inflammation via prostaglandin synthesis. The inflammation is resolved by LA metabolites.

Parent LA alone potently suppressed immune response induced by bacterial challenge in *S. exigua*. This inhibitory activity may be explained by the high levels of inhibitory activity of EpOME isomers against cellular and humoral immune responses of *S. exigua*. Among the two regioisomers, 12,13-EpOME was more potent than 9,10-EpOME. This supports the previous study [9] demonstrating the role of EpOMEs as immune suppressors. In addition, the current study showed that DiHOMEs had limited inhibitory activity against cellular immune responses. These data suggest that DiHOMEs are inactive metabolites of EpOMEs due to their catalytic hydrolysis by sEH. However, in humoral immune response, DiHOMEs actively induced selective AMP gene expression including the two attacins, defensin, gallerimycin, gloverin, lysozyme, and cecropin, which include the AMPs inhibited by EpOMEs. These data suggest that DiHOMEs are specific metabolites of EpOMEs, which prevent excessive or prolonged inhibition of immune response in insects. DiHOMEs are potent immune mediators in mammals because they are synthesized by activated neutrophils and induce chemotaxis of other neutrophils at relatively low concentrations (∼10 nM) [12, 17]. The negative controls of EpOMEs were further supported by the similar activity of their dual epoxide analog, diEpOME. Further, the two possible DiHOME analogs (THOME and THF-diol) showed limited inhibitory activity against cellular immune responses but induced AMP expression.

Compared with DiHOMEs, EpOMEs were highly cytotoxic against hemocytes of *S. exigua.* The cytotoxic activity of EpOMEs was attributed to apoptosis based on cellular blebbing and DNA fragmentation in hemocytes exposed to EpOMEs [18]. Treatment with a caspase inhibitor significantly rescued the hemocytes from the cytotoxic activity of 12,13-EpOME. Apoptosis is a form of programmed cell death that removes unwanted or damaged cells [19]. Some entomopathogens use apoptosis to inhibit host immune response by killing hemocytes, e.g. the entomopathogenic nematode, *Ovomermis sinensis* inhibits the immune response of *Helicoverpa armigera* [20]; *Serratia marcescens* and *Porphyromonas gingivalis* inhibit *Bombyx mori* [21–22]; and an ectoparasitoid wasp, *Pachycrepoideus vindemiae*, inhibits *Drosophila* [23]. Apoptosis, however, plays a crucial role in the host immune response especially during viral infection in both Lepidoptera and Diptera, in which a rapid induction of apoptosis restricts viral replication by removing infected cells [24]. Apoptosis in the midgut epithelium (known as sloughing) effectively blocks the pathogen entry into the insect hemocoel by eliminating infected cells and replacing the epithelium with regenerated cells [25]. In lepidopteran and dipteran insects, cell death is required for the release of prophenol oxidase (PPO) from specific hemocytes called oenocytoid (in *Spodoptera*) or crystal cells (in *Drosophila*) because PPO lacks the signal peptide and is released into plasma for activation by serine proteases [13, 26–27]. PGE_2_ triggers cell death via membrane receptors specific to the hemocytes of *S. exigua* [28]. However, the role of EpOMEs in cell lysis and activation of PPO remains unknown. Based on the delayed production of EpOMEs compared with the immune activator, EpOMEs may play a crucial role in eliminating infected or damaged cells during late immune response. Our current study also showed that the cell death induced by EpOMEs was attenuated by hydrolysis into less active forms, DiHOMEs, in *S. exigua*.

The immunosuppressive activity of EpOMEs was further supported by their alkoxide derivatives or agonists such as sEH inhibitors, which prevented their degradation into DiHOMEs. The alkoxides are synthetic derivatives obtained by replacing the epoxide in EpOME with a methoxy, ethoxy, propoxy or (iso)propoxy group. The alkoxy ethers cannot be hydrolyzed by she; however, as shown by the insect growth regulator methoprene, alkoxides can mimic epoxides in several biological systems. This significantly modifies the biological activities to suppress different immune responses, suggesting the critical importance of the epoxide structure in the immunosuppressive activity of 12,13-EpOME. Interestingly, A841 (= the methyl ester propoxy EpOME mimic) was the most potent among the eight alkoxides tested and even more potent than 12,13-EpOME itself. A similar screening of the oxylipin alkoxides using kidney cells was conducted to evaluate their role in protecting organs against reactive oxygen species during anticancer drug treatment using cisplatin [29]. The immunosuppressive activities of EpOMEs were evaluated using sEH inhibitors to upregulate the endogenous EpOME levels by blocking the conversion of EpOMEs to DiHOMEs. Urea-based compounds inhibiting sEH originally designed to inhibit mammalian sEH [30] inhibited invertebrate sEHs including nematodes [31] and insects [8–9]. Among six sEH inhibitors, AUDA and CUDA were highly potent, presumably by elevating endogenous EpOME levels, which led to the inhibition of immune response in the test insect. The dysregulation of the immune responses induced by the EpOME analogs or agonists might facilitate entomopathogens because the pathogens are usually eliminated by the immune responses. Insect immunity is known to exhibit efficient defense against Bt infection [32]. Our current study demonstrates that the EpOME analogs (diEpOME and A841) or agonist (AUDA) significantly enhanced the Bt virulence. This finding supports the role of EpOMEs as negative regulators of insect immunity and DiHOMEs are the their inactive metabolites.

This study demonstrates the immunosuppressive roles of EpOMEs mediated by inhibition of cellular and humoral immune responses in a lepidopteran insect. Further, the EpOMEs also induce apoptosis against immune cells. In contrast, their hydroxylated metabolites, DiHOMEs, are inactive in immunosuppression, suggesting that EpOMEs resolve the immune response induced by immune mediators following pathogen infection in insects (see Fig 7A). Although several immune mediators, such as nitric oxide (NO), biogenic monoamines, cytokines, and eicosanoids, are known to induce various immune responses [33–34], the termination or resolution of the induced immune response in insects has yet to be established. Injury in mammals leads to upregulation of innate and subsequent acquired immune responses to defend against pathogens. The immune response is later resolved at the site of injury by anti-inflammatory mediators such as resolvins, protectins, glucocorticoids, catecholamines, prostaglandins (PGs) of the E-series, NO, and interleukins [35–36]. In addition to the inhibition of cellular and humoral immune responses by EpOMEs, their high cytotoxic activities against hemocytes via apoptosis is crucial in resolving the immune responses in insects. Immune response is ultimately suppressed by the overt induction of apoptosis in immune cells endogenously, and is well known in mammals [37]. The first known mediator suppressing insect immunity was PGI_2_ in *S. exigua* [38]. Interestingly, the genes of EpOME synthase or PGI_2_ are expressed at a late stage after the immune challenge probably to terminate the induced immune response. The study findings suggest that EpOMEs play a crucial role in resolving immune response and DiHOMEs are the inactive forms in insects. More broadly, these data suggest that insects like mammals exhibit an active mechanism for resolving as well as initiating inflammation. A comparative analysis of diverse species can provide possible insight into the underlying regulatory mechanisms of initiation and resolution of inflammation.

## 4. Materials and methods

### 4.1 Insect rearing

Larvae of *S. exigua* were collected from Welsh onion (*Allium fistulosum* L.) fields in Andong, Korea. The larvae were reared under laboratory conditions (27 ± 1°C, 16:8 h (L:D), and 60 ± 5% relative humidity) with an artificial diet [39]. They underwent five instars (L1∼L5) before pupation. A diet of sugar solution (10%) was fed to adults and an empty Petri dish was provided for oviposition.

### 4.2 Chemicals

9,10-EpOME, 12,13-EpOME, 9,10-DiHOME, 12,13-DiHOME, LA, and prostaglandin E_2_ (PGE_2_) were purchased from Cayman (Ann Arbor, MI, USA). A caspase inhibitor Z-DEVD-FMK (DEVD) was purchased from Sigma-Aldrich Korea (Seoul, Korea). Other EpOME derivatives (diEpOME, THOME, and THF-diol) were prepared as described in previous studies involving the synthesis of oxygenated LA metabolites [40–41]. sEH inhibitors (AUDA, CUDA, 1661, 1663, 1659, and 1213) were prepared as reported previously for the synthesis of urea-pharmacophore-based inhibitors of the sEH enzyme [30, 42]. Four EpOME mimics (A839-A842) were generated via oxymercuration-demercuration, as previously described for the synthesis of arachidonic acid-derived epoxyeicosatrienoic acid (EET) mimics [29], using methyl LA with an appropriate alcohol as the solvent (methanol, ethanol, *n*-propanol or *iso*-propanol, respectively), followed by purification on silica gel to obtain the corresponding mono-alkoxide as a mixture of four regioisomers. The free acids (A843-A846) were obtained by basic hydrolysis of the corresponding methyl esters (A839-A842, respectively). The synthesis of A839-A846 is described in the Supporting Information (Figs. S3 and S4). An anticoagulant buffer (ACB, pH 4.5) was prepared with 186 mM of NaCl, 17 mM of Na_2_EDTA, and 41 mM of citric acid. Phosphate-buffered saline (PBS) was prepared with 100 mM phosphoric acid and the pH was adjusted to 7.4 using 1 N NaOH.

### 4.3 Hemocyte-spreading behavior assay

Two-day-old L4 larvae of *S. exigua* were used to analyze the hemocyte-spreading behavior. Approximately 250 µL of hemolymph was collected from five larvae and mixed with 750 µL of cold ACB, and incubated on ice for 20 min. The diluted hemolymph was then centrifuged at 800×*g* for 5 min at 4°C to obtain the pellet, which was re-suspended in 300 μL of filter-sterilized TC-100 insect cell culture medium (Welgene, Gyeongsan, Korea). Next, 9 μL of hemocyte suspension was incubated with 1 μL of a test chemical on a glass coverslip for 40 min at room temperature (RT). After incubation, hemocytes were fixed with 4% paraformaldehyde for 10 min at 25°C, followed by washing three times with filter-sterilized 1× PBS. Hemocytes were then permeabilized with 0.2% Triton-X in PBS for 2 min at 25°C. Hemocytes were then washed three times, followed by incubation with 5% skim milk for 10 min at 25°C and subsequently with fluorescein isothiocyanate (FITC)-tagged phalloidin in PBS for 60 min. After washing three times, the cells were incubated with 4’,6-diamidino-2-phenylindole (DAPI, 1 mg/mL). Finally, after washing twice in PBS, the cells were observed under a fluorescence microscope (DM2500, Leica, Wetzlar, Germany) at 400 × magnification. Hemocyte-spreading was determined based on the extension of F-actin growth beyond the original cell boundary of hemocytes. The cell spread was scored by counting the cells exhibiting F-actin growth in 100 randomly selected cells. Each treatment was performed in triplicate using independent hemocyte preparations.

### 4.4 RNA extraction, cDNA preparation, and qPCR

To extract RNA, the intestine was removed from the larvae to avoid any contamination due to non-target organisms. Total RNAs were extracted from the larvae using TRIzol reagent (Invitrogen, Carlsbad, CA, USA) according to the manufacturer’s instructions. The extracted RNA was used to synthesize complementary DNA (cDNA) using RT-premix (Intron Biotechnology, Seoul, Korea) containing an oligo-dT primer. The cDNA was quantified using a spectrophotometer (NanoDrop, Thermo Fisher Scientific, Wilmington, DE, USA). The cDNA (80 ng per µL) was used as a template for quantitative PCR (qPCR) using gene-specific primers (Table S1). The qPCR was performed using SYBR Green real-time PCR master mixture (Toyobo, Osaka, Japan) as described by Bustin et al. [43] using a real-time PCR system (Step One Plus Real-Time PCR System, Applied Biosystems, Singapore). The reaction mixture (20 μL) contained 10 μL of Power SYBR Green PCR Master Mix, 1 μL of cDNA template (80 ng), and 1 μL each of forward and reverse primers, and 7 µL of deionized distilled water. The qPCR was initiated by heat treatment at 95°C for 10 min, followed by 40 cycles of denaturation at 94°C for 30 s, annealing at 52°C for 30 s, and extension at 72°C for 30 s. A ribosomal protein gene, *RL32*, was used as an endogenous control. Each treatment was performed in triplicate with independent samples. Expression analysis of qPCR was performed using a comparative CT method [44].

### 4.5 AMP gene expression analysis

The levels of 10 antimicrobial peptide (AMP) genes were analyzed using L4 larvae after an immune challenge with *Escherichia coli* (1.73×10^3^ cells/larva) injection into the hemocoel. The insects were also injected with test compound(s) along with *E. coli* to assess any effect on AMP induction. Total RNA was collected as described above 8 h after treatment. After cDNA synthesis, qPCR was performed as described above using gene-specific primers (Table S1). The ribosomal protein gene, *RL32*, was amplified as the housekeeping gene serving as the internal standard in the qPCR assay. All samples were analyzed in triplicate.

### 4.6 Cellular immune assay based on hemocytic nodule formation

L4 larvae were used in bioassays to determine hemocyte nodule formation. *E. coli* was injected at a dose of 1.73×10^3^ cells per larva through the proleg and incubated at room temperature (RT) for 8 h. The insects were also injected with test compound(s) along with *E. coli* cells to assess effects on nodulation. After the dissection of test larvae, the melanized nodules were counted under a stereoscopic microscope (Stemi SV11, Zeiss, Jena, Germany) at 50× magnification. Each treatment was performed in triplicate.

### 4.7 Cytotoxicity test based on cellular blebbing

Cell blebs on the hemocyte membrane are symptoms of cytotoxicity as reported previously by Cho and Kim [45]. A test compound (1 µg per larva) was injected into L4 larvae and incubated at RT for 24 h. The hemolymph was collected into an ACB as described above. Subsequently, a hemocyte suspension (10 µL) was placed on a slide glass. The number of cell blebs were counted from 100 randomly selected cells under a phase contrast microscope (BX41, Olympus, Tokyo, Japan) at 100× magnification. Each assessment was performed in triplicate with independent insect samples.

### 4.8 Terminal deoxynucleotidyl transferase dUTP nick-end labeling (TUNEL) assay of apoptosis

A TUNEL assay to determine apoptosis was performed using an *in situ* Cell Death Detection kit (Abcam, Cambridge, UK). For this assay, 1 µg of test chemical was injected into L4 larvae and incubated for 24 h under the rearing conditions described above. After incubation, hemolymph was collected into ACB as described above. Subsequently, 10 μL of hemocyte suspension was mixed with 1 μL of 10 μM 5-bromouridine (BrdU) solution containing terminal transferase. The mixture was then placed on cover glass in a wet chamber. After 1 h incubation at RT, the medium was replaced with 4% paraformaldehyde and incubated at RT for 10 min. The cells were washed with PBS, and treated with 0.3% Triton-X in PBS, followed by incubation for 2 min at RT. After blocking with 5% bovine serum albumin in PBS for 10 min, cells were incubated with mouse anti-BrdU antibody (diluted 1:15 in blocking solution) for 1 h at RT. After removing the unbound anti-BrdU antibody, cells were incubated with FITC-conjugated anti-mouse IgG antibody (diluted 1:500 in blocking solution) for 1 h. After washing with PBS, 10 μL of DAPI was added and incubated at RT for 5 min. Cells were then washed with 10 μL of PBS. Glycerol: PBS (1:1) solution (10 μL) was then added to cells on a cover glass, which was then placed on a slide glass for observation under a fluorescence microscope (Leica) in FITC mode. Each treatment was replicated three times. Each replication represented an independent sample preparation.

### 4.9 Insecticidal bioassay

To determine the bacterial virulence against *S. exigua*, the insect pathogen, *Bacillus thuringiensis* ssp. *aizawai* (Bt), was cultured in tryptic soy broth (TSB) (Difco, Detroit, MI, USA) at 28°C. After 48 h, the cultured Bt suspension was treated at 4°C for another 48 h for sporulation. After centrifugation at 8,000 rpm for 15 min, the cell pellets were harvested and freeze-dried. Bioassay was performed using the leaf-dipping method. Briefly, a piece of cabbage leaf (2 cm^2^) was dipped into Bt suspension (10^8^ spores/mL) for 5 min. After removing the excess suspension, the treated cabbage was placed on a filter paper in a Petri dish (90 × 15 mm). Mortality was assessed at 4 days after treatment. Each treatment (= 10 larvae) was performed in triplicate.

### 4.10 Phenol oxidase activity

Phenol oxidase (PO) activity was measured using L-3,4-dihydroxyphenylalanine (DOPA) as a substrate. Each L4 larva was treated with a test chemical (1 µg) along with *E. coli* (1.73×10^3^ cells/larva). After 8 h, hemolymph was collected from the treated larva, and the plasma was separated by centrifugation at 800×*g* for 5 min at 4°C. The reaction mixture (200 µL) consisted of 10 µL of plasma, 4 µL of test compound, 10 µL of DOPA, and 176 µL of PBS. Absorbance was assessed at 495 nm using a VICTOR multi-label Plate reader (PerkinElmer, Waltham, MA, USA). The activity was expressed as change in absorbance per min per plasma (ABS/min/µL). Each treatment was replicated three times.

### 4.11 Insect sample preparation to quantify EpOMEs and DiHOMEs

Two-day-old L4 larvae of *S. exigua* were used for extraction of PGE_2_, EpOMEs, and DiHOMEs. The tissue samples were collected from the whole body without gut to avoid exogenous contamination. *E. coli* (1.73 × 10^3^ cells/larva) was injected into the insect hemocoel through the proleg. An insect body sample was collected from ∼0.85 g of larvae at different time points (0 ∼ 48 h) into a 15 mL tube and washed three times with chilled PBS. The tissue preparation was performed in triplicate. Each sample was homogenized three times (10 min per cycle) in PBS with an ultrasonicator (Bandelin Sonoplus, Berlin, Germany) at 80% power, and subsequently adjusted to a pH of 4.0 using 1 N HCl. The sample was mixed with 1 mL ethyl acetate to obtain an organic hyper-phase. The aquatic phase was extracted twice again with ethyl acetate. The combined ethyl acetate extracts containing free fatty acids or their derivatives were concentrated to ∼500 μL under a gentle nitrogen stream and applied to a small silicic acid column (2 × 90 mm containing 30 mg of Type 60A, 100–200 mesh silicic acid, Sigma-Aldrich Korea). Extracts were sequentially eluted with 250 μL of increasingly polar solvents starting with 100% ethyl acetate, followed by ethyl acetate:acetonitrile (1:1, v:v), acetonitrile:methanol (1:1, v:v), and 100% methanol. All the fractions were used to quantify PGE_2_, EpOMEs, and DiHOMEs. Each treatment was replicated three times with independent sample preparation.

### 4.12 Measurement of PGE_2_, EpOMEs, and DiHOMEs using LC-MS/MS

LC-MS/MS was conducted using a QTrap 4500 (AB Sciex, Framingham, MA, USA) equipped with an auto-sampler, a binary pump, and a column oven. The analytical column was an Osaka Soda (Osaka, Japan) C18 column (2.1 mm × 150 mm, 2.7 μm) maintained at 40°C. The mobile phases consisted of 0.1% formic acid in water and 0.1% formic acid in acetonitrile. The linear gradient was: 30% B at 0 min, 30% B at 2 min, 65% B at 12 min, 95% B at 12.5 min, 95% B at 25.0 min, 30% B at 28.0 min, and 30% B at 30 min. The flow rate was 0.40 mL/min. The auto sampler was set at 5°C and the injection volume was 10 μL. LC-MS/MS was equipped with an electrospray ionization (ESI) source. ESI was performed in negative ion mode. The source parameters after optimization were: source temperature, 600°C; curtain gas flow rate, 32 L/min; ion source gas flow rate, 60 L/min; and the adjusted spray voltage, −4,000 V. Analyses were performed in multiple reaction monitoring detection modes using nitrogen as collision gas. MassView1.1 software (AB Sciex, Seoul, Korea) were used for peak detection, integration, and quantitative analysis.

### 4.13 Statistical analysis

Continuous dependent variables were subjected to one-way analysis of variance (ANOVA) using PROC GLM in the SAS program [46]. Mortality data were transformed by arsine for normalization. All experiments were performed in three independent replicates. The means ± standard errors (SD) were plotted using the Sigma Plot (Systat Software, Point Richmond, CA, USA). The means were compared with the least significant difference (LSD) test involving a Type I error of 0.05.

## Supplementary Information

**S1 Table. Primers used in this study**

**S1 Fig. LC-MS/MS chromatograms of PGE_2_, EpOMEs, and DiHOMEs extracted from the gut-removed *S. exigua* larvae.** Each PGE_2_, EpOME and DiHOME analysis was performed using five different doses of standard.

**S2 Fig. Chemical structures of EpOME analogs and agonists.** (**A**) Structure of diEpOME, THOME, and THF-diols. (**B**) Structure of eight different EpOME alkoxides. (**C**) Structure of six sEH inhibitors.

**S3 Fig. General procedure for synthesis of EpOME-mimics A839-A842**. Methyl linoleate (4.0 mmol) was dissolved in the appropriate alcohol (25 mL, methanol, ethanol, *n*-propanol, and *iso*- propanol for the synthesis of A839, A840, A841, and A842, respectively), followed by the addition of HgOAc_2_ (6.0 mmol). The mixture was stirred under argon overnight, followed by dropwise addition of freshly prepared aqueous solution of NaBH_4_ (8.0 mmol in 8 mL water) at 0°C. After stirring for 10 min, the mixture was treated with AcOH (10 drops). Stirring was stopped to allow the precipitated Hg (l) to settle. The supernatant was transferred to a round-bottomed flask. The remaining Hg (l) was resuspended in EtOH (5 mL) twice, and the supernatants were collected and combined with the first supernatant. The combined supernatant was evaporated in vacuo, and the residue was transferred to a separating funnel with water (100 mL), followed by extraction with hexane (3 x 30 mL). The combined organic extract was washed with sat. aq. NaHCO_3_ (1 x 20 mL) and dried (MgSO_4_); the solvent was removed in vacuo. The residue was purified by flash chromatography on silica gel using a 3-5% EtOAc gradient and heptane as eluent. A regioisomeric mixture of the desired mono-alkoxides (A839, A840, A841, or A842 was obtained as a colorless oil in 35-46% chemical yield.

**(A) A839**: ^1^H NMR (400 MHz, CDCl_3_) δ 5.49-5.29 (m, 2H), 3,66 (s, 3H), 3.31-3.29 (m, 3H), 3.20-3.09 (m, 1H), 2.33-2.15 (m, 3H), 2.10-1.94 (m, 3H), 1.65-1.22 (m, 20H), 0.91-0.84 (m, 3H). HRMS (ESI): Exact mass calculated for C_20_H_38_NaO_3_: 349.2713, found 349.2713.

**(B) A840**: ^1^H NMR (400 MHz, CDCl_3_) δ 5.48-5.30 (m, 2H), 3,66 (s, 3H), 3.59-3.39 (m, 2H), 3.27-3.16 (m, 1H), 2.33-2.15 (m, 3H), 2.10-1.96 (m, 3H), 1.65-1.23 (m, 20H), 1.20-1.14 (m, 3H), 0.91-0.85 (m, 3H). HRMS (ESI): Exact mass calculated for C_21_H_40_NaO_3_: 363.2870, found 363.2869.

**(C) A841**: ^1^H NMR (400 MHz, CDCl_3_) δ 5.49-5.29 (m, 2H), 3,66 (s, 3H), 3.47-3.14 (m, 3H), 2.33-1.94 (m, 6H), 1.65-1.22 (m, 22H), 0.97-0.83 (m, 6H). HRMS (ESI): Exact mass calculated for C_22_H_42_NaO_3_: 377.3026, found 377.3025.

**(D) A842**: ^1^H NMR (400 MHz, CDCl_3_) δ 5.48-5.20 (m, 2H), 3.65 (s, 3H), 3.62-3.55 (m, 1H), 3.32-3.21 (m, 1H), 2.34-1.97 (m, 6H), 1.67-1.55 (m, 2H), 1.50-1.20 (m, 18H), 1.17-1.09 (m, 6H), 0.92.0.84 (m, 6H). HRMS (ESI): Exact mass calculated for C_22_H_42_NaO_3_: 377.3026, found 377.3025.

**S4 Fig. General procedure for synthesis of EpOME-mimics A843-A846**. The corresponding methyl ester (200 mg A839, A840, A841 or A842, respectively) was dissolved in a 1:1 mixture of THF/MeOH (20 mL). A solution of LiOH (30 equiv.) in water (5 mL) was then added, and the mixture was stirred under argon for 16 h at ambient temperature. The reaction mixture was quenched by the addition of aq. HCl (40 mL, 1,0 M) and extracted with EtOAc (2 x 20 mL). The combined organic extract was washed with water (2 x 10 mL), dried (MgSO_4_) and evaporated. The residue was purified by passing it through a short plug of silica gel using a gradient of 10-30% EtOAc in heptane to afford the desired carboxylic acid (A843, A844, A845 or A846) in 77-88% yield as a colorless oil.

**(A) A843**: ^1^H NMR (400 MHz, CDCl_3_) δ 11.46-9.20 (bs, 1H), 5.50-5.28 (m, 2H), 3.36-3.29 (m, 3H), 3.20-3.10 (m, 1H), 2.33 (t, 2H, *J* = 7,6 Hz), 2.29-2.15 (m, 1H), 2.10-1.94 (m, 3H), 1.70-1.57 (m, 2H), 1.56-1.12 (m, 18H), 0.91-0.82 (m, 3H). HRMS (ESI): Exact mass calculated for C_19_H_35_O_3_: 311.2592, found 311.2591.

**(B) A844**: ^1^H NMR (400 MHz, CDCl_3_) δ 11.37-8.85 (bs, 1H), 5.49-5.21 (m, 2H), 3.60-3.39 (m, 2H), 3.28-3.17 (m, 1H), 2.33 (t, 2H, *J* = 7,6 Hz), 2.28-2.15 (m, 1H), 2.11-1.93 (m, 3H), 1.67-1.57 (m, 2H), 1.55-1.21 (m, 18H), 1.21-1.14 (m, 3H), 0.91-0.83 (m, 3H).HRMS (ESI): Exact mass calculated for C_20_H_37_O_3_: 325.2748, found 325.2748.

**(C) A845**: ^1^H NMR (400 MHz, CDCl_3_) δ 11.78-8.55 (bs, 1H), 5.49-5.28 (m, 2H), 3.49-3.28 (m, 2H), 3.27-3.16 (m, 1H), 2.33 (t, 2H, *J* = 7.5 Hz), 2.29-2.14 (m, 1H), 2.12-1.93 (m, 3H), 1.67-1.21 (m, 22H), 0.95-9.82 (m, 6H). HRMS (EI): Exact mass calculated for C_21_H_39_O_3_: 339.2905, found 339.2904.

**(D) A846**: ^1^H NMR (400 MHz, CDCl_3_) δ 12.15-8.92 (bs, 1H), 5.50-5.27 (m, 2H), 3.69-3.54 (m, 1H), 3.33-3.20 (m, 1H), 2.34 (t, 2H, *J* = 7,5 Hz), 2.19 (t, 1H, J = 6,2 Hz), 2.11-1.93 (m, 3H), 1.68-1.55 (m, 2H), 1.50-1.19 (m, 18H), 1.17-1.07 (m, 6H), 0.91-0.81 (m, 3H). HRMS (EI): Exact mass calculated for C_21_H_39_O_3_: 339.2905, found 339.2904.

## Data Availability Statement

The authors confirm that all data underlying the findings are fully available without restriction. All relevant data are within the paper and its Supporting Information file.

## Author Contributions

**Conceptualization**: Bruce D. Hammock, Yonggyun Kim.

**Data curation**: Md Tafim Hossain Hrithrik, Dong-Hee Lee, Nalin Singh, Anders Vik.

**Formal analysis**: Md Tafim Hossain Hrithrik, Dong-Hee Lee.

**Funding acquisition**: Anders Vik, Bruce D. Hammock, Yonggyun Kim.

**Investigation**: Md Tafim Hossain Hrithrik, Dong-Hee Lee, Nalin Singh, Anders Vik, Bruce D. Hammock, Yonggyun Kim.

**Methodology**: Md Tafim Hossain Hrithrik, Dong-Hee Lee, Nalin Singh, Anders Vik.

**Project administration**: Anders Vik, Bruce D. Hammock, Yonggyun Kim. **Resources**: Nalin Singh, Anders Vik, Bruce D. Hammock, Yonggyun Kim. **Software**: Md Tafim Hossain Hrithrik, Dong-Hee Lee.

**Supervision**: Anders Vik, Bruce D. Hammock, Yonggyun Kim.

**Validation**: Md Tafim Hossain Hrithrik, Dong-Hee Lee, Nalin Singh, Anders Vik.

**Visualization**: Md Tafim Hossain Hrithrik, Dong-Hee Lee, Anders Vik.

**Writing – original draft**: Md Tafim Hossain Hrithrik, Dong-Hee Lee, Nalin Singh, Anders Vik, Bruce D. Hammock, Yonggyun Kim.

**Writing – review & editing**: Nalin Singh, Anders Vik, Bruce D. Hammock, Yonggyun Kim.

